# Real-time Plasmid Transmission Detection Pipeline

**DOI:** 10.1101/2024.07.09.602722

**Authors:** Natalie Scherff, Jörg Rothgänger, Thomas Weniger, Alexander Mellmann, Dag Harmsen

## Abstract

The spread of antimicrobial resistance among bacteria by horizontal plasmid transmissions poses a major challenge for clinical microbiology. Here, we evaluate a new real-time plasmid transmission detection pipeline implemented in the SeqSphere^+^ (Ridom GmbH, Münster, Germany) software.

Within the pipeline, a local Mash plasmid database is created and Mash searches with a distance threshold of 0.001 are used to trigger plasmid transmission early warning alerts (EWA). Clonal transmissions are detected using cgMLST allelic differences. The integrated tools MOB-suite, NCBI AMRFinderPlus, CGE MobileElementFinder, pyGenomeViz, and MUMmer are used to characterize plasmids and for visual pairwise plasmid comparisons, respectively. We evaluated the pipeline using published hybrid assemblies (Oxford Nanopore Technology/Illumina) of a surveillance and outbreak dataset with plasmid transmissions. To emulate prospective usage, samples were imported in chronological order of sampling date. Different combinations of the user-adjustable parameters sketch size (1,000 vs 10,000) and plasmid size correction were tested and discrepancies between resulting clusters were analyzed with Quast.

When using a sketch size of 1,000 with size correction turned on, the SeqSphere^+^ pipeline agreed with the published data and produced the same clonal and carbapenemase-carrying plasmid clusters. EWAs were in the correct chronological order.

In summary, the developed pipeline presented here is suitable for integration into clinical microbiology settings with limited bioinformatics knowledge due to its automated analyses and alert system, which are combined with the GUI-based SeqSphere^+^ platform. Thus, with its integrated sample database, (near) real-time plasmid transmission detection is within reach in bacterial routine-diagnostic settings when long-read sequencing is employed.

**Importance:** Plasmid-mediated spread of antimicrobial resistance (AMR) is a major challenge for clinical microbiology and monitoring of potential plasmid transmissions is essential to combat further dissemination. Whole-genome sequencing (WGS) is often used to surveil nosocomial transmissions but usually limited to the detection of clonal transmissions (based on chromosomal markers). Recent advances in long-read sequencing technologies enable full reconstruction of plasmids and the detection of very similar plasmids but so far easy-to-use bioinformatic tools for this purpose were missing. Here we present an evaluation of an innovative real-time plasmid transmission detection pipeline. It is integrated into the GUI-based SeqSphere^+^ software, which already offers cgMLST based pathogen outbreak detection. It requires very limited bioinformatics knowledge, and its database, automated analyses, and alert system make it well suited for prospective clinical application.

## Introduction

Antimicrobial resistance (AMR) is a major challenge in clinical microbiology as it is limiting the therapeutic options to manage infectious diseases (1). This is largely mediated by the rapid spread of resistance genes among bacterial populations via horizontal gene transfer (HGT) facilitated by mobile genetic elements (MGE) such as plasmids (2). Plasmids are extrachromosomal DNA elements that can serve as a vehicle to exchange genes between different bacterial strains and even species (3).

Recently, multi-species hospital outbreaks of carbapenem-resistant bacteria have been reported (4, 5). Whole-genome sequencing (WGS) based surveillance has become an important tool to monitor nosocomial transmissions in many hospitals (6), but these activities are usually limited to the detection of clonal transmissions, i.e., based on chromosomal markers, of single species. Thus, tools are needed that enable the detection of similar plasmids from WGS data. A variety of software tools has been developed for plasmid reconstruction and characterization (reviewed in (7)). Among the most popular are currently CGE PlasmidFinder (8), COPLA (9), PLACNET (10), and MOB-suite (11), which relies on Mash (12) for comparing k-mers of plasmids. All these tools require some form of bioinformatic expertise, which is often lacking in routine microbiology laboratories. Another major limitation is that plasmids are generally difficult to fully reconstruct from short-reads due to their high number of repetitive regions (7). However, with the recent advances in long-read sequencing, e.g., Oxford Nanopore Technology (ONT), accurate long-read data will become more available to clinical microbiology laboratories (13).

The GUI-based Ridom SeqSphere^+^ software (14) is widely used in clinical microbiology laboratories. One of its main purposes is the detection of clonal transmissions using core-genome multi-locus sequence typing (cgMLST) based similarity of bacterial isolates. Here, we evaluated the newly implemented real-time plasmid detection pipeline to enable users to detect potential plasmid transmissions within their own WGS datasets complementary to clonal transmissions. We used two already published datasets of hybrid assemblies of carbapenem-resistant bacteria from a United Kingdom (UK) (15) and a United States (US) hospital (16), respectively.

## Materials and Methods

### SeqSphere^+^ real-time plasmid transmission detection pipeline

We developed the pipeline as an extra module for the SeqSphere^+^ (Ridom GmbH, Münster, Germany) software. The first steps involve the reconstruction and characterization of plasmids from assembled data. These steps are based on the MOB-suite (v3.18) software package (11) and work for both short- and long-read data. Short-read and non-circular long read data go into the MOB-recon module first to identify and reconstruct plasmids. The reconstructed plasmids are then forwarded to MOB-typer, which assigns a primary and secondary cluster ID (Mash distance cluster threshold = 0.006 and 0.025, respectively), rep and relaxase type(s), and mobility prediction to each plasmid. For long-read data, the FASTA headers are checked for circularity terms ([topology = circular]) and only non-circular contigs a forwarded to MOB-recon. Circular contigs that are above 500kb in size are considered to be chromosomes, smaller contigs are considered to be plasmids and directly go into MOB-typer. Optionally, the MOB-recon module can also be skipped using a [non-recon] term in the FASTA header. AMR targets are determined using the NCBI AMRFinderPlus (v3.11.26) (17). Integrated mobile genetic elements (iMGEs) are detected with CGE MobileElementFinder (v1.1.2) (18). The typing results from MOB-typer, AMRFinderPlus, and MobileElementFinder are then summarized in a tabular “Chromosome & Plasmid” overview. From here, plasmid contigs can be exported as FASTA files for downstream analyses.

AMRFinderPlus results are grouped into two columns, one of them only showing priority AMR genes, which are defined as targets that might confer resistance to carbapenems, colistin, vancomycin, or methicillin, and genes that contain ESBL or AmpC in their name. The detected iMGE(s) are also shown in a separate column if they enclose a priority AMR gene.

### Long-read data plasmid transmission analysis module

With the recently introduced version 10 of the SeqSphere^+^ software with the long-read data plasmid transmission analysis module, potential plasmid transmissions can be detected using Mash (v2.1) comparisons (12). The identified plasmids can be used to build a local Mash plasmid database from the users’ own data. Pairwise Mash distances can then be utilized to detect very similar plasmids. This information can be used in several ways. First, if new plasmids are imported via the pipeline mode, they are compared with all existing plasmids in the database and if there are matches below a certain threshold (default = 0.001), an early warning alert (EWA) is triggered and shown on the SeqSphere^+^ start screen. No EWA is generated if the respective samples contain a cgMLST scheme and have a clonal (i.e., cgMLST allelic distance below the species-specific threshold) relationship. Second, the complete plasmid database can be used to perform an all-against-all comparison to create an exportable distance matrix and/or a single-linkage clustering. Third, the database can be searched for similar plasmids from a single plasmid of interest. This plasmid search can be triggered from both the “Chromosome & Plasmid” overview and EWA reports. Within the resulting table, sample metadata and plasmid typing results for each plasmid are included. Finally, partially annotated plasmids (origin(s) of replication, priority AMR gene(s), other AMR gene(s), and iMGE(s)) can also be pairwise aligned and visualized using MUMmer v3.23 (19) and pyGenomeViz (v0.4.4; https://moshi4.github.io/pyGenomeViz/). To create a better visualization, there is an optional fixstart function to change the start and orientation of contigs. It uses *dnaA* for chromosomes and for plasmids the first occurrence of a *rep* gene as the start of the reoriented contig. Figure 1 shows an overview of the complete pipeline.

**Figure 1:**
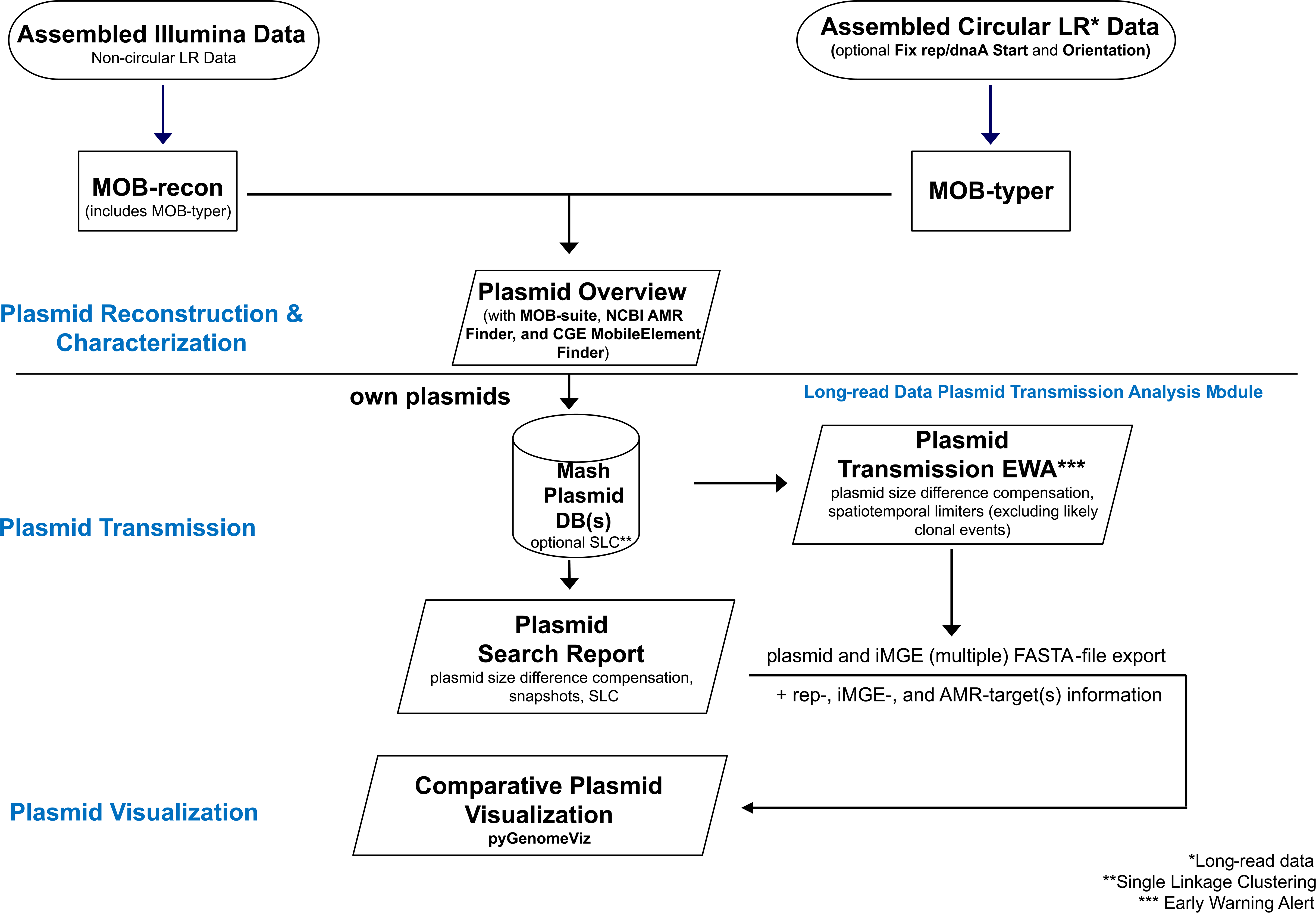
Overview of plasmid transmission detection pipeline.

SeqSphere^+^ allows for configuration of multiple parameters including sketch size, sequence length compensation, and Mash distance threshold for matches (default:0.001). For the sketch size the user has the choice between 1,000 and 10,000. The k-mer size of 21 is fixed. Due to the large variability in plasmid length and as Mash distances favor contigs of similar size we have implemented a scaling factor for comparing plasmids with difference in unique content where the match threshold is lowered for each 1% of size difference. The default is a lowering of 0.0003 per 1% size difference up to a limit of 40% size difference but both the lowering value and the upper limit can be adjusted by the user.

### Evaluation of the pipeline using two published datasets

To test the pipeline functionality and evaluate parameter settings, the pipeline was applied to two different published datasets. The first was a set of 85 isolates of phenotypically carbapenem-resistant Gram-negative bacteria from a single hospital in the UK (Addenbrooke’s Hospital, Cambridge) that were collected over a period of six years (Addenbrookes dataset) (15). We excluded four samples without a detected carbapenemase gene leading to a total number of 81 samples in our test dataset. These isolates were originally sequenced using a combination of Illumina HiSeq and Oxford Nanopore Technologies (ONT) MinION R9.4.1 technologies. Original hybrid assemblies were created using Flye (v2.8) (20), Unicycler (v0.4.8) (21) and Canu (v2.1.1) (22). These hybrid assemblies were downloaded in FASTA format from the European Nucleotide Archive (ENA), where they had been previously deposited under the study accession PRJEB30134. A Mash search against the NCBI RefSeq genomes was performed to confirm species assignment and rule out potential contamination. From the information in the original publication, we did assume that each contig, except the chromosome, was a single plasmid. However, since we did not have circularity information, we used a [non-recon] term in the FASTA headers to skip the MOB-recon part as additional plasmid reconstruction was not done in the original publication. The study was done retrospectively but to test the EWAs, we emulated a prospective analysis by importing the files in the chronological order of sampling date. Sampling date and patient information were taken from the supplemental material provided with the original publication.

Plasmid clusters were defined as plasmids with a Mash distance ≤ 0.001. In addition to clusters, the number of potential transmissions was counted. If a sample carried multiple plasmids that belonged to different plasmid clusters, it was assessed whether all of these plasmids were transferred to the same or different receiving samples. If all plasmids were shared with the same sample, this was counted as one co-transmission event. If plasmids of one sample were connected to different samples, each connection was counted as a single plasmid transmission event. If two samples had a clonal relation, the respective plasmids were not counted as a plasmid but as a clonal transmission event.

To assess clonal relations, we used cgMLST allelic distances. For species with public cgMLST schemes hosted on the Ridom nomenclature server (www.cgMLST.org), we used the species-specific default threshold. For species without a public scheme, we created *ad hoc* schemes, using the NCBI Genomes reference sequence of each species as seed genome and a threshold of 15 alleles to determine clonal clusters. To check for additional transmissions of only iMGEs, a user-defined task template was created from an allele database that contained all iMGE with a carbapenemase gene that were found using the CGE MobileElementFinder with thresholds of ≥90% identity and ≥95% alignment length.

For parameter evaluation, we tested four combinations: a sketch size of 1,000 and 10,000, each with and without size compensation applied to the Mash distance. A cluster threshold of ≤ 0.001 was used for all four approaches. Plasmid clusters with discrepancies between these different runs were analyzed with Quast (v5.2) (23). Here, the longest plasmid of each cluster was set as reference. Mismatches, InDels, and larger gaps were analyzed and counted. . Finally, the pipeline was applied to the non carbapenemase-carrying plasmids of this dataset as well.

To prove that the pipeline with the determined Mash parameters also works in another setting with different strains, we evaluated a second dataset. This one contained 19 isolates of a multispecies (n = 7) outbreak of *bla*NDM-5-producing Enterobacterales in a US hospital (UPMC Presbyterian dataset) (16). The isolates were collected from February 2021 to February 2023 from 15 patients. All isolates were sequenced on an Illumina NextSeq 550 and ONT MinION with R9.4.1 flow cells. Hybrid assemblies were created using Unicycer v0.5.0 and deposited by the original authors for download at NCBI under BioProject PRJNA981541. In the original paper, outbreak plasmids were defined as *IncX3* plasmids harboring *bla*NDM-5, “that had sequence coverage values (*i.e.*, the proportion of each plasmid’s sequence that was found in the Illumina genome) of at least 95%, and a sequence identity of <15 single nucleotide polymorphisms (SNPs) per 100 kb of plasmid-sequence compared with the fully resolved, circular plasmid sequence” (16). For our evaluation, the same analysis as described above was applied to the carbapenemase-carrying plasmids of this dataset.

## Results

### Analysis of clonal clusters in the Addenbrookes dataset

Clonal clusters were determined based on cgMLST allelic distances with species-specific thresholds (15 alleles difference for species without public cgMLST schemes). In total, seven clonal clusters were detected, three for *Klebsiella pneumoniae,* two for *Escherichia coli,* one for *Pseudomonas putida*, and one for *Enterobacter hormaechei* (Table 1). Except one *K. pneumoniae* cluster comprising eleven isolates (kleb_2), all clusters contained two isolates. The allelic distances were between 0 and 11. In addition, we found two *E. roggenkampii* isolates with a distance of one allele but since these isolates were from the same patient, they were denoted as intra-host variability.

**Table 1:** Overview of detected clonal and carbapenemase-carrying plasmid transmission clusters. Cluster names correspond to the original publication.

| Cluster | No. of isolates | Species | Carbapenemase gene | Rep types |
| --- | --- | --- | --- | --- |
| **Clonal clusters** |  |  |  |  |
| pseu_1 | 2 | *Pseudomonas putida* | IMP-70 |  |
| ecoli_1 | 2 | *Escherichia coli* | OXA-232 |  |
| ecoli_2 | 2 | *Escherichia coli* | NDM-5 |  |
| ehor_1 | 2 | *Enterobacter hormaechei* | IMP-70 |  |
| kleb_1 | 2 | *Klebsiella pneumoniae* | NDM-1 |  |
| kleb_2 | 11 | *Klebsiella pneumoniae* | NDM-1 |  |
| kleb_3 | 2 | *Klebsiella pneumoniae* | OXA-232 |  |
| **Intra-host variability** |  |  |  |  |
| erog_1 | 2 | *Enterobacter roggenkampii* | OXA-181 |  |
| **Plasmid clusters** |  |  |  |  |
| Clust_33 | 2 | *Escherichia coli, Klebsiella pneumoniae* | NDM-6 | IncFIB |
| Clust_133 | 2 | *Acinetobacter baumannii* | OXA-72 | rep_1127 |
| Clust_17 | 4 | *Escherichia coli, Klebsiella aerogenes, Klebsiella pneumonia* | OXA-48 | IncI |
| Clust_148* | 4 | *Escherichia coli, Klebsiella pneumoniae* | NDM-1 | IncC |
| Clust_54 | 2 | *Escherichia coli* | NDM-5 | IncFIB/IncFIA |
| Clust_9* | 3 | *Enterobacter hormaechei* | IMP-70 | IncC |
| Clust_72* | 8 | *Escherichia coli, Klebsiella pneumoniae* | OXA-232 | Col |
| Clust_98 | 6 | *Escherichia coli, Enterobacter hormaechei, Klebsiella pneumonia* | OXA-181 | IncX3 |
| Clust_217 | 4 | *Enterobacter roggenkampii, Klebsiella grimontii, Raoultella ornithinolytica* | OXA-181 | IncX3 |

\* includes clonal isolates

### Analysis of carbapenemase-carrying plasmids in the Addenbrookes dataset

Initially, we only analyzed the plasmids carrying a carbapenemase gene. Out of the 81 isolates, eleven isolates contained chromosomally encoded carbapenemase genes. The other 70 isolates had carbapenemase-carrying plasmids. One isolate comprised two carbapenemase-carrying plasmids leading to a total number of 71 analyzed plasmids.

By re-sequencing strains from documented and published plasmid transmission events with ONT and Pacific Biosciences sequencing methodology (4, 24), we determined a Mash distance of 0.001 as suitable for recovering nearly identical plasmids (data not shown). By varying the unique content of those plasmids, we came up with a scaling factor of 0.0003 per 1% plasmid size difference. Next, using a sketch size of 10,000, a Mash distance threshold of 0.001, and a 0.0003 size compensation, we found nine plasmid clusters (Table 1). One additional plasmid cluster consisted of only clonal isolates (kleb_2). Three plasmid clusters (Clust_148, Clust_9, Clust_72) partly contained clonal isolates. The clusters comprised between two and eight isolates from one to three different species. The relatedness of the 71 carbapenemase-carrying plasmids is visualized with a population snapshot using the exported clustering distance matrix (Figure 2).

**Figure 2:**
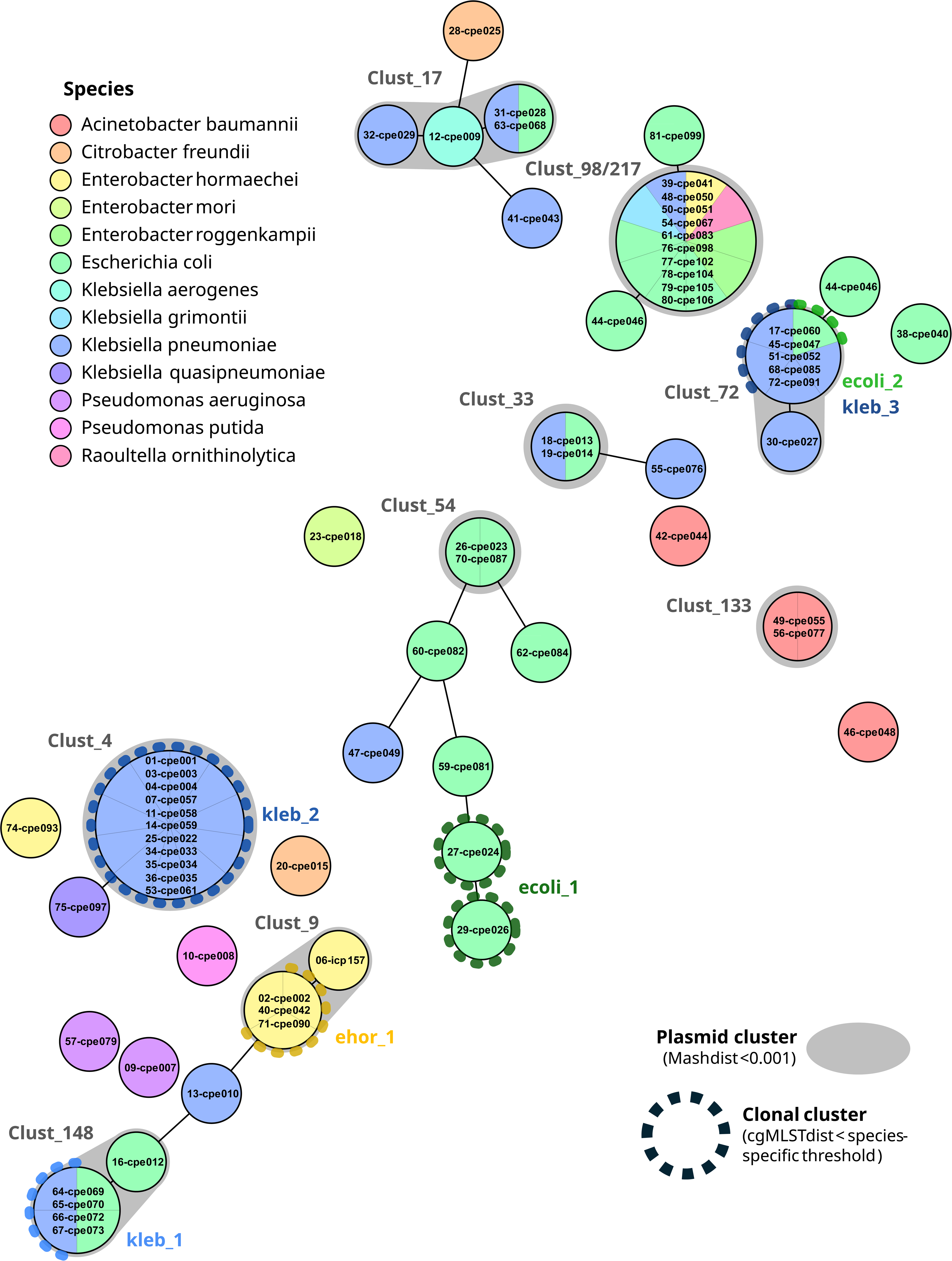
Plasmid population snapshot based on Mash distances between 71 carbapenemase-carrying plasmids. Each circle represents one plasmid, plasmids with a Mash distance of 0 are lumped together in single circles with a larger size. Connection lines are drawn for plasmids with a Mash distance <0.06. Fill color represents the species of the respective isolates and colored, dotted circles mark clonal clusters among these isolates. Plasmid clusters (Mash distance threshold <0.001) are illustrated by grey shaded areas around the circles. Clonal (in color) and plasmid (in grey) cluster names are stated next to the clusters.

To evaluate the Mash parameters, we repeated this analysis with a lower sketch size of 1,000 and tested both sketch sizes with and without size compensation and compared the resulting clusters. Out of the nine plasmid clusters, four showed discrepancies between the different methods (Table 2). One plasmid pair (Clust_54) was missed with a sketch size of 1,000 but only when size compensation was turned off. In Clust_9, one additional sample was included in the cluster for both sketch sizes when size compensation was applied. Clust_148 also contained an additional plasmid with size compensation, but only with a sketch size of 10,000. Finally, Clust_98/Clust_217 was lumped together and contained nine samples with size compensation but without size compensation it was divided in two clusters with six and three samples, respectively. We counted the number of differences between the disputable plasmid and the reference plasmid of each cluster.

**Table 2:** Plasmids with variations in clustering results between different Mash database parameters. (Compensated) Mash distances and Quast results for each comparison of a plasmid with a reference plasmid (grey fill color) are shown. The different parameters were sketch size of 1,000 (1k) or 10,000 (10k) and use of size compensation (sc). Mash distance values that led to different clustering results (threshold = 0.001) are highlighted in orange. The dashed line marks two clusters that were joined into one cluster with size correction but separated without.

| Plasmid clusters |  |  | Mash distance to reference |  |  |  | Quast results |  |  |  |
| --- | --- | --- | --- | --- | --- | --- | --- | --- | --- | --- |
| Cluster | Plasmid size (kb) | Size difference to reference % | 1k | 1k + sc | 10k | 10k + sc | Mismatches | Indels | Large gaps | Sum events |
| Clust_9 | 127,2 | 0% |  |  |  |  |  |  |  |  |
|  | 127,2 | 0% | 0,0000 | 0,0000 | 0,0000 | 0,0000 | 0 | 0 | 0 | 0 |
|  | 126,2 | 1% | 0,0002 | 0,0000 | 0,0000 | 0,0000 | 0 | 1 | 2 | 3 |
|  | 96,3 | 24% | 0,0068 | 0,0000 | 0,0074 | 0,0002 | 1 | 10 | 2 | 13 |
| Clust_148 | 145,8 | 0% |  |  |  |  |  |  |  |  |
|  | 145,8 | 0% | 0,0000 | 0,0000 | 0,0000 | 0,0000 | 0 | 0 | 0 | 0 |
|  | 145,8 | 0% | 0,0000 | 0,0000 | 0,0000 | 0,0000 | 0 | 0 | 0 | 0 |
|  | 140,1 | 4% | 0,0022 | 0,00103 | 0,0019 | 0,0007 | 4 | 0 | 1 | 5 |
| Clust_54 | 153,2 | 0% |  |  |  |  |  |  |  |  |
|  | 148,4 | 3% | 0,00101 | 0,0001 | 0,00099 | 0,00004 | 1 | 1 | 1 | 3 |
| Clust_98 | 51,5 | 0% | 0,0000 | 0,0000 | 0,0000 | 0,0000 |  |  |  |  |
|  | 51,5 | 0% | 0,0000 | 0,0000 | 0,0000 | 0,0000 |  |  |  |  |
|  | 51,5 | 0% | 0,0000 | 0,0000 | 0,0000 | 0,0000 |  |  |  |  |
|  | 51,5 | 0% | 0,0000 | 0,0000 | 0,0000 | 0,0000 |  |  |  |  |
|  | 51,5 | 0% | 0,0000 | 0,0000 | 0,0000 | 0,0000 |  |  |  |  |
|  | 51,5 | 0% |  |  |  |  |  |  |  |  |
| Clust_217 | 47,1 | 9% | 0,0021 | 0,0000 | 0,0022 | 0 | 1 | 0 | 1 | 2 |
|  | 47,1 | 9% | 0,0021 | 0,0000 | 0,0022 | 0 | 1 | 0 | 1 | 2 |
|  | 48,5 | 6% | 0,0022 | 0,0004 | 0,0022 | 0,0005 | 1 | 0 | 1 | 2 |

For Clust_54 there were three, for Clust_148 five, and for Clust_9 13 differences, respectively. For Clust_98/Clust_217 we compared all samples of Clust_217 with one reference of Clust_98, which resulted in two differences.

In one cluster (Clust_72), we detected a case, where one plasmid (size = 12,282 bp) contained a duplicated sequence, which was otherwise completely identical with another plasmid (size = 6,141 bp) that only contained the single sequence, i.e. their uncorrected Mash distance was 0. We checked for additional transmissions of iMGEs by retrieving the sequences of all iMGEs containing a carbapenemase gene and searching for these sequences in all isolates utilizing a user-defined task template. No additional transmissions of iMGEs were found using this approach.

### Analysis of all plasmids in the Addenbrookes dataset

After analysis of the carbapenemase-carrying plasmids, in contrast to the original publication, we also had a look at the overall plasmid landscape of these samples. In total, 324 plasmids were found in 78 of the 81 samples (three samples did not contain plasmids). Each sample carried 1-10 plasmids with an average of four plasmids per sample. Of these, 175 plasmids from 58 samples belonged to a cluster (Mash distance 0.001). On average, three plasmids per isolate belonged to a cluster with more than one plasmid. A total of 51 clusters (including clonal and carbapenemase-carrying plasmid clusters) was detected with 2-11 isolates per cluster (median = 2, mean = 3.4).

In addition to clonal and plasmid clusters, we counted the number of potential transmission events. In total, 44 single transmission events were counted. Of these, twelve were clonal transmissions, 18 were single plasmid transmissions, and 14 were co-transmissions of two or more plasmids to the same sample. Three of the single and five of the co-plasmid-transmissions were intra-host transmissions. Fifteen isolates transferred more than one plasmid; for nine of them, all plasmids were transferred to the same and for six of them to different receiving samples. Furthermore, we detected two cases where an isolate received plasmids from a clonal and an additional plasmid transmission. One of these involved two different species, *K. pneumoniae* and *E. coli* (Figure 3). Here, the two *K. pneumoniae* samples had identical MLST sequence types (STs) and cgMLST allelic profiles. However, the sample that was isolated later carried additional plasmids. Two of these plasmids (size = 1.8 kb & 5.2 kb) clustered with plasmids of an *E. coli* isolate that also shared a 145 kb *bla*NDM1-carrying plasmid with both *K. pneumoniae* samples.

**Figure 3:**
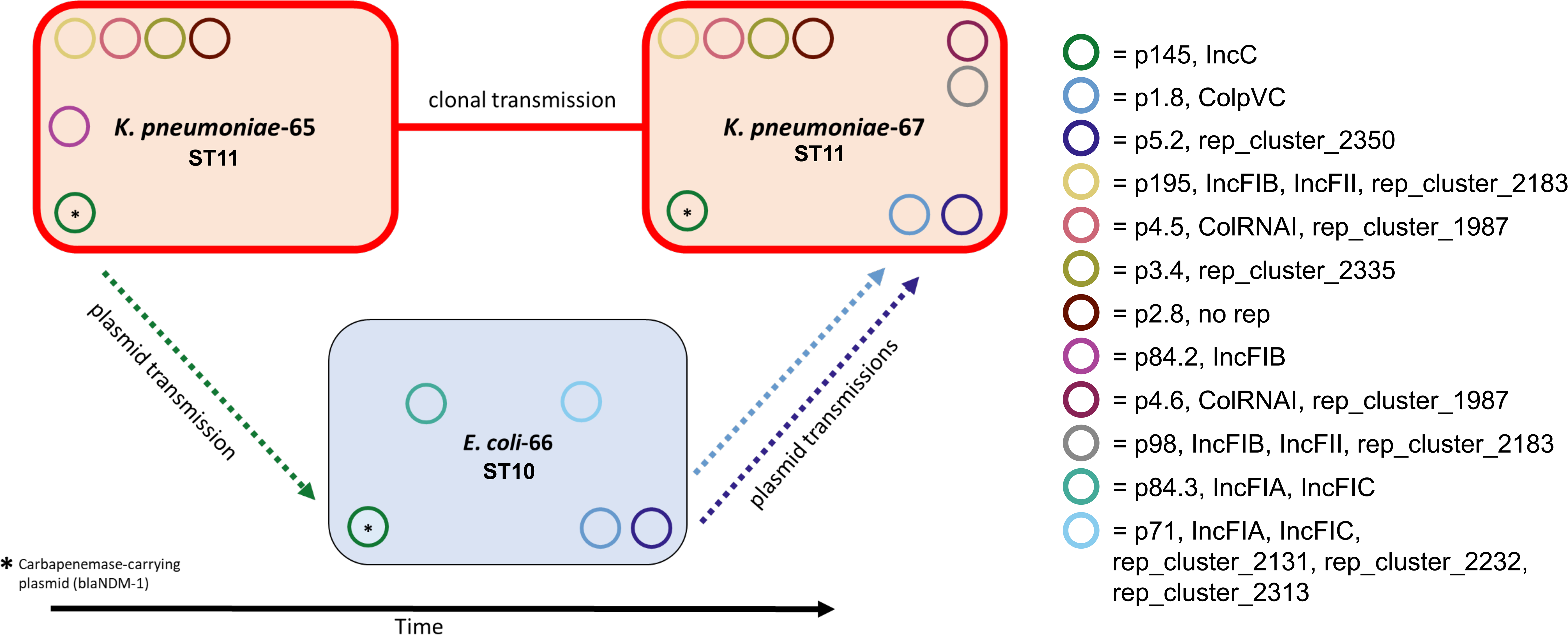
Visualization of a combination of clonal and horizontal plasmid transmissions. Direction of transmission is assumed based on temporal order of isolation. ST = MLST sequence type.

### Analysis of carbapenemase-carrying plasmids in the UPMC Presbyterian dataset

As a further evaluation, we analyzed the carbapenemase-carrying plasmids of a second dataset, comprising 19 isolates of a multispecies *bla*NDM-5 outbreak (16). According to the original publication, all isolates belonged to a single outbreak, where *bla*NDM-5 was encoded on a 46.2 kb *IncX3* family plasmid. The plasmids differed by only 0-2 SNPs with the exception of one shorter (45.2 kb) and one larger (55.8 kb) plasmid that lost or acquired a transposase gene, respectively.

Using our pipeline with a sketch size of 1,000 and size correction turned on, all 19 plasmids were put into a single plasmid cluster. However, when size correction was turned off, the larger (17% size difference) plasmid did not cluster with the remaining plasmids (uncorrected Mash distance = 0.003). Both, *bla*NDM-5 and *IncX3* were detected on all plasmids by AMRFinderPlus and MOB-suite, respectively. In addition, among other information, relaxase type MOBP was assigned by MOB-typer and the plasmids were predicted to be conjugative.

## Discussion

In this study, we used two published datasets of plasmids from carbapenem-resistant bacteria to evaluate a new plasmid detection pipeline implemented in the SeqSphere^+^ software. We determined a Mash distance of 0.001 and a scaling factor of 0.0003 per 1% plasmid size difference as suitable for recovering nearly identical plasmids. Thresholds like these are always somewhat arbitrary and need to be a compromise between sensitivity and specificity. However, the developers of the MOB-suite came up with a distance threshold of 0.025 for short-read sequencing data (25). With long-read data this value can be lower. Independent from our own considerations, Roberts *et al.* came up with the same threshold of 0.001 (15). For both datasets, our results are in complete agreement with the findings from the original studies, in particular, the same clonal and plasmid clusters were found. Moreover, the same resistance genes and plasmid rep-types were detected, despite using different tools and databases. In the Roberts *et al*. study, Abricate (https://github.com/tseemann/abricate) was used with the CARD (26) and PlasmidFinder (8) databases, while our pipeline is based on NCBI AMRFinderPlus (17) and MOB-suite (11). In both the Roberts *et al.* and our study, Mash with a sketch size of 10,000 and a distance threshold of 0.001 was used to detect plasmid results, leading not surprisingly to the same cluster results. However, in contrast to a file-based Mash approach, our automated, database-based pipeline enables a prospective real-time surveillance of plasmid transmissions.

Raabe *et al.* defined outbreak plasmids based on Illumina sequence coverage values of at least 95%, and a sequence identity of <15 SNPs per 100 kb of plasmid sequence when compared against the complete reference plasmid sequence. They also used the CGE PlasmidFinder for rep-typing. Different approaches were also used for clonal cluster detection. While Roberts *et al.* used a combination of SNPs and split k-mer analysis (SKA), the SeqSphere^+^ pipeline is based on cgMLST allelic distances.

We compared the clustering results of Mash databases constructed with a sketch size of 1,000 and 10,000. Both are values used for plasmids in the literature (15, 25). While the higher sketch size increases the sensitivity, it also requires more computational memory and power and enlarges the database size. We found that a sketch size of 1,000 was generally enough to detect the same clusters at a threshold of 0.001, with one exception. The sensitivity can be increased without a significant increase in computational power requirements by applying a size compensation. The size compensation allows to detect potential transmissions of plasmids that share the same backbone but may have lost or gained larger unique segments. This was shown with the UPMC Presbyterian dataset, where one insertion of a transposase gene increased the size of the otherwise identical plasmid by 17 %. The Mash distance between this larger plasmid and the other outbreak plasmids was above the clustering threshold without size correction. However, size compensation must have an upper limit to prevent too many false positives just resulting from shared promiscuous iMGEs. An interesting finding was a plasmid pair, where one plasmid consisted of two exact copies of another plasmid. This is likely a result of multimer formation. To account for such phenomena, we decided to limit the size compensation to a maximum of 40% plasmid size differences but keep the uncorrected Mash distance value for plasmids with a larger size difference instead of ignoring plasmid pairs above 40% size difference. Thus, plasmids with large size differences but very similar content, as for example multimers, can still be detected.

When analyzing the whole plasmid landscape of the Addenbrookes dataset, we found that the reconstruction of potential transmission scenarios can be complicated due to complex co-transmission patterns. Based on genetic clustering, the number of plasmid transmission events within the analyzed dataset is estimated to be twice as common as clonal transmission of pathogens but detailed epidemiological information is needed to confirm these results. In particular, it is still largely unknown how often very similar plasmids occur without a transmission, i.e. as “core plasmids” of certain lineages or species. As other plasmid comparison tools, the pipeline only works with well reconstructed, ideally circular plasmids and thus requires long-read sequences. Moreover, Mash as a k-mer-based approach does not take synteny into account. Although same content is usually also same synteny in a local context, it is possible to check for rearrangements between plasmid sequences by utilizing pyGenomeViz, which is implemented in SeqSphere+, as a visual approach. For more diverse data, alternatively, a new tool such as Pling (27), which is alignment-based and thereby more compute-intensive, could be used. Currently the literature is still lacking analyses of plasmid transmissions based on long-read data. However, with long-read sequencing becoming more and more achievable for clinical laboratories, the need for suitable analysis tools is growing. We presented here a retrospective re-analysis of published datasets. However, in a prospective surveillance scenario, every single transmission would be captured by the EWA system as soon as the isolates are imported, allowing for a near real-time plasmid transmission detection.

In conclusion, the new SeqSphere^+^ plasmid transmission detection pipeline allows the detection of plasmid clusters in both retrospective (single linkage clustering of plasmid databases) and prospective (Early Warning Alerts) studies. The automated, database-based approach enables near real-time surveillance and allows for rapid informed actions.

## Acknowledgments

We thank Leah Roberts (CIIC, Queensland University of Technology) for providing help with sequence and epidemiological data. Further, we thank James Robertson and Kyrylo Bessonov (both National Microbiology Laboratory, Public Health Agency of Canada) for critical reading and improvement of the manuscript.

## Disclaimers

NS, JR, and TW are (part-time) employees of Ridom GmbH. JR, TW, and DH are shareholders of Ridom GmbH. AM declares no conflict of interest.

## References

1. Murray CJ, Ikuta KS, Sharara F, Swetschinski L, Aguilar GR, Gray A, Han C, Bisignano C, Rao P, Wool E, Johnson SC, Browne AJ, Chipeta MG, Fell F, Hackett S, Haines-Woodhouse G, Hamadani BHK, Kumaran EAP, McManigal B, Agarwal R, Akech S, Albertson S, Amuasi J, Andrews J, Aravkin A, Ashley E, Bailey F, Baker S, Basnyat B, Bekker A, Bender R, Bethou A, Bielicki J, Boonkasidecha S, Bukosia J, Carvalheiro C, Castañeda-Orjuela C, Chansamouth V, Chaurasia S, Chiurchiù S, Chowdhury F, Cook AJ, Cooper B, Cressey TR, Criollo-Mora E, Cunningham M, Darboe S, Day NPJ, Luca MD, Dokova K, Dramowski A, Dunachie SJ, Eckmanns T, Eibach D, Emami A, Feasey N, Fisher-Pearson N, Forrest K, Garrett D, Gastmeier P, Giref AZ, Greer RC, Gupta V, Haller S, Haselbeck A, Hay SI, Holm M, Hopkins S, Iregbu KC, Jacobs J, Jarovsky D, Javanmardi F, Khorana M, Kissoon N, Kobeissi E, Kostyanev T, Krapp F, Krumkamp R, Kumar A, Kyu HH, Lim C, Limmathurotsakul D, Loftus MJ, Lunn M, Ma J, Mturi N, Munera-Huertas T, Musicha P, Mussi-Pinhata MM, Nakamura T, Nanavati R, Nangia S, Newton P, Ngoun C, Novotney A, Nwakanma D, Obiero CW, Olivas-Martinez A, Olliaro P, Ooko E, Ortiz-Brizuela E, Peleg AY, Perrone C, Plakkal N, Ponce-de-Leon A, Raad M, Ramdin T, Riddell A, Roberts T, Robotham JV, Roca A, Rudd KE, Russell N, Schnall J, Scott JAG, Shivamallappa M, Sifuentes-Osornio J, Steenkeste N, Stewardson AJ, Stoeva T, Tasak N, Thaiprakong A, Thwaites G, Turner C, Turner P, Doorn HR van, Velaphi S, Vongpradith A, Vu H, Walsh T, Waner S, Wangrangsimakul T, Wozniak T, Zheng P, Sartorius B, Lopez AD, Stergachis A, Moore C, Dolecek C, Naghavi M. 2022. Global burden of bacterial antimicrobial resistance in 2019: a systematic analysis. The Lancet 399:629–655.

2. Sheppard AE, Stoesser N, Wilson DJ, Sebra R, Kasarskis A, Anson LW, Giess A, Pankhurst LJ, Vaughan A, Grim CJ, Cox HL, Yeh AJ, Modernising Medical Microbiology (MMM) Informatics Group, Sifri CD, Walker AS, Peto TE, Crook DW, Mathers AJ. 2016. Nested Russian doll-like genetic mobility drives rapid dissemination of the carbapenem resistance gene *bla*KPC. Antimicrob Agents Chemother 60:3767–3778.

3. Orlek A, Stoesser N, Anjum MF, Doumith M, Ellington MJ, Peto T, Crook D, Woodford N, Sarah Walker A, Phan H, Sheppard AE. 2017. Plasmid classification in an era of whole-genome sequencing: Application in studies of antibiotic resistance epidemiology. Frontiers in Microbiology 8:1–10.

4. Weber RE, Pietsch M, Frühauf A, Pfeifer Y, Martin M, Luft D, Gatermann S, Pfennigwerth N, Kaase M, Werner G, Fuchs S. 2019. IS26-mediated transfer of *bla*NDM–1 as the main route of resistance transmission during a polyclonal, multispecies outbreak in a German hospital. Frontiers in Microbiology 10:1–14.

5. Marimuthu K, Venkatachalam I, Koh V, Harbarth S, Perencevich E, Cherng BPZ, Fong RKC, Pada SK, Ooi ST, Smitasin N, Thoon KC, Tambyah PA, Hsu LY, Koh TH, De PP, Tan TY, Chan D, Deepak RN, Tee NWS, Kwa A, Cai Y, Teo Y-Y, Thevasagayam NM, Prakki SRS, Xu W, Khong WX, Henderson D, Stoesser N, Eyre DW, Crook D, Ang M, Lin RTP, Chow A, Cook AR, Teo J, Ng OT, Carbapenemase-Producing Enterobacteriaceae in Singapore (CaPES) Study Group, Marimuthu K, Venkatachalam I, Cherng BPZ, Fong RKC, Pada SK, Ooi ST, Smitasin N, Thoon KC, Hsu LY, Koh TH, De PP, Tan TY, Chan D, Deepak RN, Tee NWS, Ang M, Lin RTP, Teo J, Ng OT. 2022. Whole genome sequencing reveals hidden transmission of carbapenemase-producing Enterobacterales. Nat Commun 13:3052.

6. Mellmann A, Bletz S, Böking T, Kipp F, Becker K, Schultes A, Prior K, Harmsen D. 2016. Real-time genome sequencing of resistant bacteria provides precision infection control in an institutional setting. Journal of Clinical Microbiology 54:2874–2881.

7. Paganini JA, Plantinga NL, Arredondo-Alonso S, Willems RJL, Schürch AC. 2021. Recovering *Escherichia coli* plasmids in the absence of long-read sequencing data. Microorganisms 9:1613.

8. Carattoli A, Zankari E, Garcia-Fernandez A, Voldby Larsen M, Lund O, Villa L, Moller Aarestrup F, Hasman H. 2014. *In silico* detection and typing of plasmids using PlasmidFinder and plasmid multilocus sequence typing. Antimicrobial agents and chemotherapy 58:3895–3903.

9. Redondo-Salvo S, Bartomeus-Peñalver R, Vielva L, Tagg KA, Webb HE, Fernández-López R, de la Cruz F. 2021. COPLA, a taxonomic classifier of plasmids. BMC Bioinformatics 22:390.

10. Vielva L, de Toro M, Lanza VF, de la Cruz F. 2017. PLACNETw: a web-based tool for plasmid reconstruction from bacterial genomes. Bioinformatics 33:3796–3798.

11. Robertson J, Nash JHE. 2018. MOB-suite: software tools for clustering, reconstruction and typing of plasmids from draft assemblies. Microbial Genomics 4.

12. Ondov BD, Treangen TJ, Melsted P, Mallonee AB, Bergman NH, Koren S, Phillippy AM. 2016. Mash: Fast genome and metagenome distance estimation using MinHash. Genome Biology 17:1–14.

13. Sereika M, Kirkegaard RH, Karst SM, Michaelsen TY, Sørensen EA, Wollenberg RD, Albertsen M. 2022. Oxford Nanopore R10.4 long-read sequencing enables the generation of near-finished bacterial genomes from pure cultures and metagenomes without short-read or reference polishing. Nat Methods 19:823–826.

14. Jünemann S, Sedlazeck FJ, Prior K, Albersmeier A, John U, Kalinowski J, Mellmann A, Goesmann A, von Haeseler A, Stoye J, Harmsen D. 2013. Updating benchtop sequencing performance comparison. Nature biotechnology 31:294–296.

15. Roberts LW, Enoch DA, Khokhar F, Blackwell GA, Wilson H, Warne B, Gouliouris T, Iqbal Z, Török ME. 2023. Long-read sequencing reveals genomic diversity and associated plasmid movement of carbapenemase-producing bacteria in a UK hospital over 6 years. Microbial Genomics 9:001048.

16. Raabe NJ, Valek AL, Griffith MP, Mills E, Waggle K, Srinivasa VR, Ayres AM, Bradford C, Creager HM, Pless LL, Sundermann AJ, Van Tyne D, Snyder GM, Harrison LH. 2024. Real-time genomic epidemiologic investigation of a multispecies plasmid-associated hospital outbreak of NDM-5-producing Enterobacterales infections. International Journal of Infectious Diseases 142:106971.

17. Feldgarden M, Brover V, Gonzalez-Escalona N, Frye JG, Haendiges J, Haft DH, Hoffmann M, Pettengill JB, Prasad AB, Tillman GE, Tyson GH, Klimke W. 2021. AMRFinderPlus and the Reference Gene Catalog facilitate examination of the genomic links among antimicrobial resistance, stress response, and virulence. Scientific Reports 11:1–9.

18. Johansson MHK, Bortolaia V, Tansirichaiya S, Aarestrup FM, Roberts AP, Petersen TN. 2021. Detection of mobile genetic elements associated with antibiotic resistance in *Salmonella enterica* using a newly developed web tool: MobileElementFinder. J Antimicrob Chemother 76:101–109.

19. Kurtz S, Phillippy A, Delcher AL, Smoot M, Shumway M, Antonescu C, Salzberg SL. 2004. Versatile and open software for comparing large genomes. Genome Biol 5:R12.

20. Kolmogorov M, Yuan J, Lin Y, Pevzner PA. 2019. Assembly of long, error-prone reads using repeat graphs. Nat Biotechnol 37:540–546.

21. Wick RR, Judd LM, Gorrie CL, Holt KE. 2017. Unicycler: Resolving bacterial genome assemblies from short and long sequencing reads. PLoS Comput Biol 13:e1005595.

22. Koren S, Walenz BP, Berlin K, Miller JR, Bergman NH, Phillippy AM. 2017. Canu: scalable and accurate long-read assembly via adaptive k-mer weighting and repeat separation. Genome Res 27:722–736.

23. Mikheenko A, Prjibelski A, Saveliev V, Antipov D, Gurevich A. 2018. Versatile genome assembly evaluation with QUAST-LG. Bioinformatics 34:i142–i150.

24. van Almsick V, Schuler F, Mellmann A, Schwierzeck V. 2022. The use of long-read sequencing technologies in infection control: Horizontal transfer of a *bla*CTX-M-27 containing lncFII plasmid in a patient screening sample. Microorganisms 10:491.

25. Robertson J, Bessonov K, Schonfeld J, Nash JHE. 2020. Universal whole-sequence-based plasmid typing and its utility to prediction of host range and epidemiological surveillance. Microbial Genomics 6:1–12.

26. Jia B, Raphenya AR, Alcock B, Waglechner N, Guo P, Tsang KK, Lago BA, Dave BM, Pereira S, Sharma AN, Doshi S, Courtot MM, Lo R, Williams LE, Frye JG, Elsayegh T, Sardar D, Westman EL, Pawlowski AC, Johnson TA, Brinkman FSLL, Wright GD, McArthur AG. 2017. CARD 2017: Expansion and model-centric curation of the comprehensive antibiotic resistance database. Nucleic Acids Research 45:D566–D573.

27. Frolova D, Lima L, Roberts L, Bohnenkämper L, Wittler R, Stoye J, Iqbal Z. 2024. Applying rearrangement distances to enable plasmid epidemiology with pling. bioRxiv 10.1101/2024.06.12.598623.

